# Beyond pseudotime: Following T-cell maturation in single-cell RNAseq time series

**DOI:** 10.1101/219188

**Authors:** David S. Fischer, Anna K. Fiedler, Eric Kernfeld, Ryan M. J. Genga, Jan Hasenauer, Rene Maehr, Fabian J. Theis

## Abstract

Cellular development has traditionally been described as a series of transitions between discrete cell states, such as the sequence of double negative, double positive and single positive stages in T-cell development. Recent advances in single cell transcriptomics suggest an alternative description of development, in which cells follow continuous transcriptomic trajectories. A cell’s state along such a trajectory can be captured with pseudotemporal ordering, which however is not able to predict development of the system in real time. We present pseudodynamics, a mathematical framework that integrates time-series and genetic knock-out information with such transcriptome-based descriptions in order to describe and analyze the real-time evolution of the system. Pseudodynamics models the distribution of a cell population across a continuous cell state coordinate over time based on a stochastic differential equation along developmental trajectories and random switching between trajectories in branching regions. To illustrate feasibility, we use pseudodynamics to estimate cell-state-dependent growth and differentiation of thymic T-cell development. The model approximates a developmental potential function (Waddington’s landscape) and suggests that thymic T-cell development is biphasic and not strictly deterministic before beta-selection. Pseudodynamics generalizes classical discrete population models to continuous states and thus opens possibilities such as probabilistic model selection to single cell genomics.

## Introductory paragraph

Single-cell experiments, such as single-cell-RNA-seq^1^, ‐qPCR^2^ and mass cytometry^3^, have recently enabled researchers to study the heterogeneity of cell populations. It was found in many studies that development proceeds in an asynchronous fashion in populations^4,5^. Therefore, pseudotemporal ordering methods have been devised to describe development as a transition in transcriptomic state rather than a transition in real time^4,5^. These pseudotemporal orderings have been leveraged in many cases to give detailed gene expression profiles along developmental processes with much higher resolution than it would have been possible based on the sampling time coordinate^6^. However, pseudotemporal orderings are static descriptions which do not include dynamic information which can be obtained from time series experiments or steady state systems for example. Steady-state problems were previously addressed in the context of cell cycle transitions^7,8^ based on discretized population structures. The problem of developmental trajectory estimation from time series data is not a steady-state problem which was recently addressed via an optimal transport framework for discrete transitions^9^ (Fig. 1b). Here, we present pseudodynamics (Fig. 1a), a mathematical framework tailored to developmental trajectories which accounts both for the continuous population structure and the non-steady state dynamics of the system to understand population growth and differentiation characteristics along trajectories. As an example, we establish an unbiased view of T-cell development based on a branching pseudotemporal ordering of cells observed with single-cell RNA-seq data and validate it with previous findings. Secondly, we show that the population growth rate can be fit as a transcriptomic state dependent function which maps out selection pressure during T-cell maturation on specific transcriptomic states. Thirdly, we show how pseudodynamics facilitates the integration of wildtype and mutant data to annotate developmental trajectories with developmental checkpoints using the example of developmental arrest of T-cells at beta-selection in Rag1 and Rag2 knock-out mice. Our model extends previous efforts on modelling gene expression distributions in time^10^ by population size dynamics and by the notion of developmental trajectories in transcriptome space. Pseudodynamics is independent of the method with which the pseudotemporal ordering is generated. In summary, pseudodynamics adds the following layers of information to a lineage trajectory: (1) population growth dynamics such as population bursts and selection, (2) an approximation of the developmental potential function including stability information of cell states, (3) an exact mapping of developmental checkpoints on a trajectory given mutant data, (4) imputation of the population density in cell state at a missing time point for experiment planning, and (5) model selection between multiple dynamic models such as identification of regions of diffusive and deterministic dynamics. We made the pseudodynamics code accessible via GitHub (https://github.com/theislab/pseudodynamics) and built an interactive website that allows exploration of the trajectory description of the T-cell development data set presented here (*not yet public*).

A population of cells observed in single-cell RNA-seq experiment is a probability density on the transcriptome space. We describe a cell’s trajectory, for example across a developmental time course, by this density as a function of time in a time series experiment (Fig. 1a), non-normalized in order to allow for population size changes. The size of the population at a given time point is the integral of the density with respect to cell state. The underlying cell state space is high-dimensional which impedes the estimation of underlying dynamic model. We therefore reduce the cell state space to a one dimensional pseudotime coordinate that measures developmental progress from a root progenitor cell. We model branching processes as trees consisting of a set of developmental trajectories which overlap or are connected at branching regions (Fig. 1a,b, online methods). Pseudodynamics describes the time dependence of the population density in pseudotime based on a population balance model (Fig. 1c): a diffusion-advection model with an additional population growth term (online methods). We define the model parameters (diffusion, drift, birth-death rate) as spline functions of the cell state to be able to infer cell state-specific characteristics of the dynamics of the population as a maximum likelihood estimation problem. The drift parameter quantifies the directed component of development and can be interpreted as the slope of the underlying developmental potential function (“Waddington’s landscape”). The diffusion parameter quantifies the undirected component of development which accounts for stochasticity of the developmental process. Note that one can use drift and diffusion parameters to stratify a trajectory into domains of directed (drift-dominated) and undirected (diffusion-dominated) development. The birth-death parameter describes population growth through cell division and cell death. We estimate parameters of the underlying partial differential equation using a regularized spline interpolation and maximum likelihood estimation; the probabilistic description of the system thus allows for uncertainty analysis on the inferred parameters. The input to this parameter estimation consists of empirical cumulative density functions of cell state observations (cells) by sample and population size estimates by time point. We achieved numerical stability and accuracy of the forward simulations necessary for parameter estimation on the partial differential equation system with the finite volumes method (online methods).

As an example, we fit the pseudodynamics model to a single-cell RNA-seq data set of mouse embryonic stem cell differentiation^1^, in which developmental progression can be captured as a pseudotemporal ordering along a single trajectory in a diffusion map^4^ (Fig. 1c, Supp. methods). Pseudodynamics was able to fit the dynamics of the population of cells along this transcriptomic trajectory and thereby extends the temporal snapshots to a continuous description in time (Fig. 1d,e, Supp. data 2). The continuous description of the population in time allows imputation of unobserved time points which we validated with leave-one-time-point-out cross validation (Fig. 1e, online methods).

Next, we applied the pseudodynamics model to thymic T-cell development observed with single-cell RNA-seq using the Drop-seq protocol^1^ in mice as 19 samples at eight different time points from 12.5 to 19.5 days after fertilisation (E12.5-P0) [cite accompanying paper from Kernfeld et al.] (Fig. 2a). We selected T-cells and putative natural killer cells in silico based on their transcriptomes (online methods). The diffusion map^11^ of this gated data set has one branching region between the T-cell lineaege and between a lineage which has high expression of natural killer cell markers (putative natural killer cells) (Fig. 2c). Indeed, T-cells and natural killer cells were previously shown to derive from the same progenitor population in the thymus^12,13^. We used diffusion pseudotime as a one dimensional cell state coordinate with the tip cell of the progenitor branch as a root cell. The cell state therefore captures transcriptomic progression along the T-cell and the putative natural killer cell lineages. We performed a linear partition of the manifold of observed cell states in diffusion component space to distinguish the T-cell and the putative natural killer cell trajectories (Supp. Fig. 1). The expression profiles along the T-cell lineage recapitulate the previously established sequence of developmental stages from double negative to double positive cells which are defined based on surface marker proteins^14^ (Fig. 2b). Moreover, the previously described sequential upregulation of transcription factors along the T-cell lineage is resolved in detail^14^ (Supp. Fig. 5). We performed a separate alignment of reads to a synthetic genome of rearranged T-cell receptor (TCR) genes to distinguish αβ‐ and γδ-T-cells. TCRα and TCRβ expression together with double-positive stage markers (Id3, Rorc, Cd4, Cd8a Cd8b1) expression suggest that the T-cell lineage trajectory in the diffusion map corresponds to the αβ-T-cell lineage (Fig. 2b). We also found TCRγ and TCRδ expressing cells on this T-cell lineage before the upregulation of double-positive stage markers (Fig. 2b) which could correspond to γδ-T-cells or to temporary expression of TCRγ and TCRδ on the αβ-T-cell lineage. The expression profiles along the T-cell lineage trajectory offer higher resolution than expression observations at previously used discrete cell stages (double negative, double-positive and single-positive stages), illustrating the order of activity of gene regulatory modules of interest (Supp. Fig. 5). Moreover, expression profiles along the trajectory can be used to suggest putative surface marker proteins for particular developmental stages (Supp. Fig. 3).

We fit pseudodynamics to the developmental tree with a single branching region between the T-cell and putative natural killer cell lineages (online methods). We extended the continuous cell state description of T-cell development by annotating the cell states with the parameter fits of the pseudodynamics model. T-cells that pass beta-selection divide rapidly and then undergo positive and negative selection^14^. The cell state-specific division and death rates are captured as a high birth-death rate after the putative point of beta-selection which then monotonically decreases with cell state and eventually becomes negative (Fig. 2d, also captured in a fit based on monocle2^5^ pseudotime: Supp. Fig. 7,8). The developmental progression throughout the double positive stage is captured as a positive drift parameter. Transcriptome-derived M‐ and G2.M-phase scores (online methods) are increased in the cell states on which positive and negative selection act, which suggests that there is an expansion of the cells that survive selection and thus add to the result that the population size is overall decreasing (negative birth-death parameter, Fig. 2e). The fit of the drift parameter along the the T-cell lineage (Fig. 2d) uncovers two intervals of fast transcriptomic development (high drift parameter) which peak at cell states 0.13 and 0.35 and one interval with drift close to zero around cell state 0.23. The two intervals of fast development correspond to transcriptomic states in which developmental transcription factors are sequentially regulated (for example Notch1 and Notch3 in interval one and Id3 and Rorc in interval two), which lead to directed transcriptomic development and to deterministic behaviour of individual cells. Indeed, we observed global changes in transcription factor activity at these stages (Fig. 2b, Supp. Fig. 5). Such intervals of directed transcriptomic development with high drift parameters correspond to regions in Waddington’s landscape with a high derivative with respect to cell state. The interval with very low drift corresponds to a region in saddle point of the landscape. We argue based on Bcl-xL^14^ and double-positive stage marker up-regulation at this cell state (Fig. 2b) that the saddle point corresponds to the transcriptomic state at the developmental check-point beta-selection. The gene expression profiles along the trajectory also suggests a regulatory mechanism for beta-selection. Developing T-cells require Bcl-xL to overcome beta-selection^14^, one of five anti-apoptotic proteins of the Bcl2 family. Indeed, Bcl-xL is upregulated after the putative beta-selection point but Bcl2 and Mcl1 are downregulated (Supp. Fig. 4c), which results in an overall Bcl2 family activity, which may no longer prevent apoptosis. The diffusion parameter is not decreased around the saddle point (Fig. 2d) which suggests that cells develop in an undirected fashion (“diffusion-dominated”) around this saddle point.

To validate the prediction of the cell state of beta-selection, we sampled Rag1 and Rag2 knockout mice which produce T-cells that cannot overcome beta-selection as they cannot rearrange the T-cell receptor genes^15^. We fit a diffusion map to the union of wild type T-cell single-cell RNA-seq samples of all timepoints, a Rag2KO at E14.5 and two Rag1KO samples at E16.5 (Supp. Fig. 9). The T-cell populations in the knock-out mice are delayed in transcriptomic development along the αβ-T-cell trajectory compared to age-matched wild type samples (Fig. 3a). The delay is statistically significant if tested with a one-sided Kolmogorov–Smirnov test on the empirical cumulative density functions across cell state of the union of mutant samples against the union of wild type samples at each time point (E14.5 p=1e-7.7, E16.5 p=1e-9.6, Supp. Fig. 10). A developmental arrest at a checkpoint such as beta-selection can cause a delay in the progression in cell state. Moreover, the absence of high cell state outliers at E16.5 which we observed in the wild type data can also be explained with a developmental arrest (Fig. 3a). Beyond trends in cell state space, we also observed a reduced mean expression of double-positive stage markers in the knock-out compared to age-matched wild-type mice (Supp. Fig. 11). We adapted the pseudodynamics model which we trained on wild type populations to account for the arrest expected in beta-selection to determine the transcriptomic state at which beta-selection occurs (online methods). We then computed a likelihood profile (Supp. methods) of the mutant data given our adapted model over cell state coordinates (Fig. 3b). The resulting maximum-likelihood estimator of the beta-selection point at cell state 0.27 is in agreement with the Bcl-xL expression profile and which marks the end of the saddle point region identified based on the drift parameter fit. We obtain similar beta-selection estimates for multiple regularization parameters (Supp. Fig. 17a).

Lastly, we evaluated the ability of pseudodynamics to impute missing time points by fitting the model on the data from all but one time point and by evaluating the fit on this withheld time point (Supp. Fig. 13-15). We found that pseudodynamics is in principle able to perform such imputation but has limited predictive power if the distributional shape changes strongly at a time point (E14.5, E17.5). Indeed one would assume that there is high uncertainty in the distribution estimate at these time points which could be uncovered with a Bayesian estimation scheme.

Developmental trajectories in cell state space are powerful descriptions which make information from very large high dimensional data sets accessible, such as through gene expression profiles across pseudotime. Temporal sample coordinates have however been difficult to exploit in such descriptions. We showed that a description solely based on transcriptomic data uncovers many known aspects of thymic T-cell development in an unbiased fashion. We showed how one can incorporate temporal information via a dynamic framework, pseudodynamics, to extend and complement the trajectory-based description. In particular, we showed that T-cell maturation is biphasic (phase 1, “towards-beta-selection”, and phase 2, “away-from-beta-selection”, Fig. 3c) which stands in contrast to the previous model which was defined based on surface markers (double-negative, double-positive and single-positive stages). One may think of this class of dynamic models as a step towards approximating the developmental potential previously termed “Waddington’s landscape”^16^ in a quantitative fashion. Indeed, one can interpret the drift parameter of the pseudodynamics model as an approximation of the negative derivative of Waddington’s landscape with respect to cell state. Accordingly, one can infer the shape of Waddington’s landscape based on the pseudodynamics fits which shows the plateau before beta-selection (Fig. 3b). This approximation of Waddington’s landscape is a quantitative description of the the degree of determinism of development of an individual cell which is related to plasticity of cellular states. We find a region of mostly indeterministic development (“diffusion-dominated”) before beta-selection. The inclusion of population size dynamics into the diffusion-drift framework allowed us to map out selection pressure and population expansion on the cell state coordinate which was not addressed in previous dynamic models for cellular development in transcriptome space^10,17^. We chose a pseudotemporal ordering as the developmental progression space in this study. Possible extensions could be using different and higher dimensional cell state spaces with coordinates such as individual marker genes or components of a dimension reduction technique, as well as ordering coordinates determined by genetic barcoding/lineage tracing^18,19^. In summary, pseudodynamics bridges the concepts of pseudotemporal ordering and cell state dynamics in a probabilistic framework that adds layers of information with uncertainty quantification to a developmental lineage.

**Figure 1.**
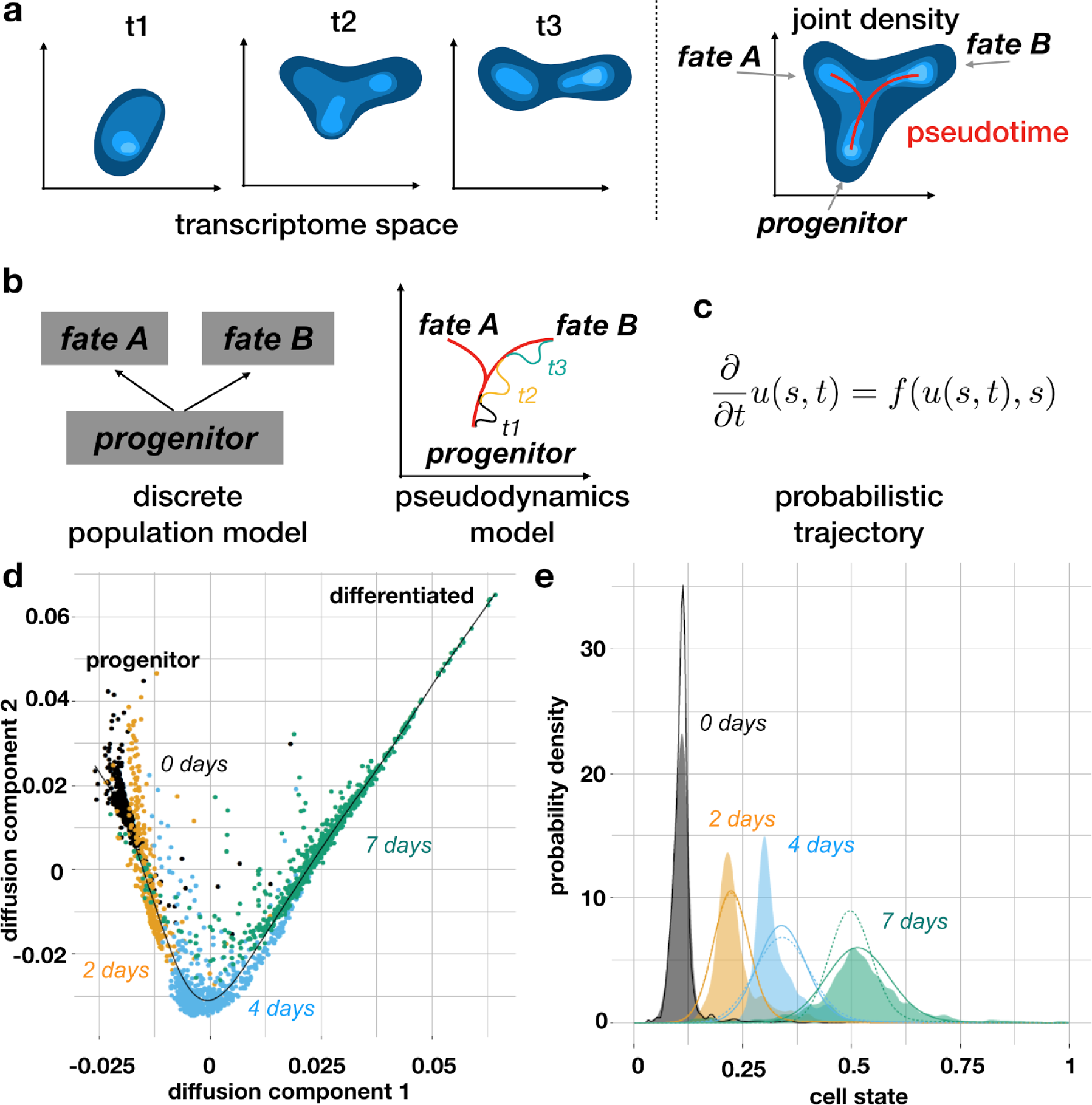
A population-based view on single-cell RNA-seq time-series experiments: Concept of pseudodynamics and example fits on a mouse embryonic stem cell differentiation data set. (**a**) Development can be modelled as a progression of a population density in transcriptome (cell state) space. Here, the developmental process is a branched lineage from a progenitor to two terminal fates. Cell state can be quantified with pseudotime. (**b**) Development described as a succession of discrete cell states versus the pseudodynamics model. (**c**) Structure of the pseudodynamics model: A trajectory in development can be defined probabilistically as change of the density *u* of cells across cell state *s* over time *t* (left hand side). This change can be modelled as a function of the density itself and cell state-specific parameters (right hand side) (online methods). (**d**) Diffusion map of mouse embryonic stem cell development in vitro after LIF removal. Colour: days after leukemia inhibitory factor (LIF) removal in cell culture. (**e**) Kernel density estimate and simulated density of cells across cell state coordinate (diffusion pseudotime) at four sampled time points. Coloured density: kernel density estimate, solid line: simulated density, dotted line (2, 4, 7 days): simulated density in leave-one-out cross-validation.

**Figure 2.**
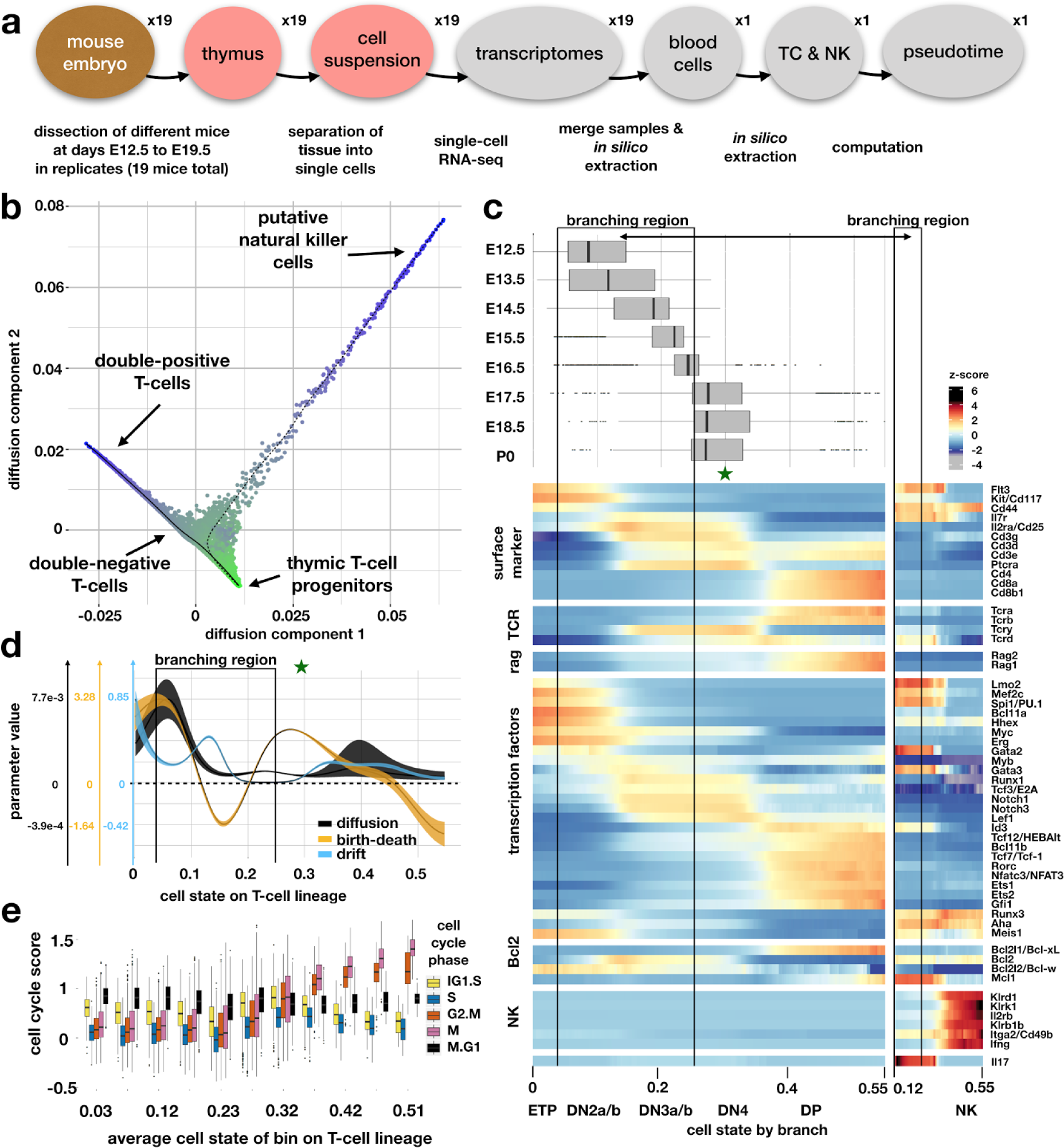
Pseudodynamics captures marker gene signatures of T-cell development in the embryonic thymus and maps out selection pressure on double-positive αβ-T-cells. (**a**) Experimental design. TC: T-cells, NK: putative natural killer cells. (**b**) Diffusion map based on TC and NK only. (**c**) Heatmap of z-scores of sliding window expression estimates across cell state (diffusion pseudotime). On top: boxplot of density of cells across cell state by time point. Red star: Mean cell state coordinate of samples in of E17.5, E18.5, P0. (**d**) Parameter estimates of pseudodynamics as spline fits of the cell state on the αβ-T-cells lineage with confidence intervals. Shaded area: spline fit to 99% confidence interval boundary on spline nodes. Green star: Mean cell state coordinate of samples of E17.5, E18.5, P0. (**e**) Transcriptome-based cell cycle state scores (online methods) per cell by cell state bin.

**Figure 3.**
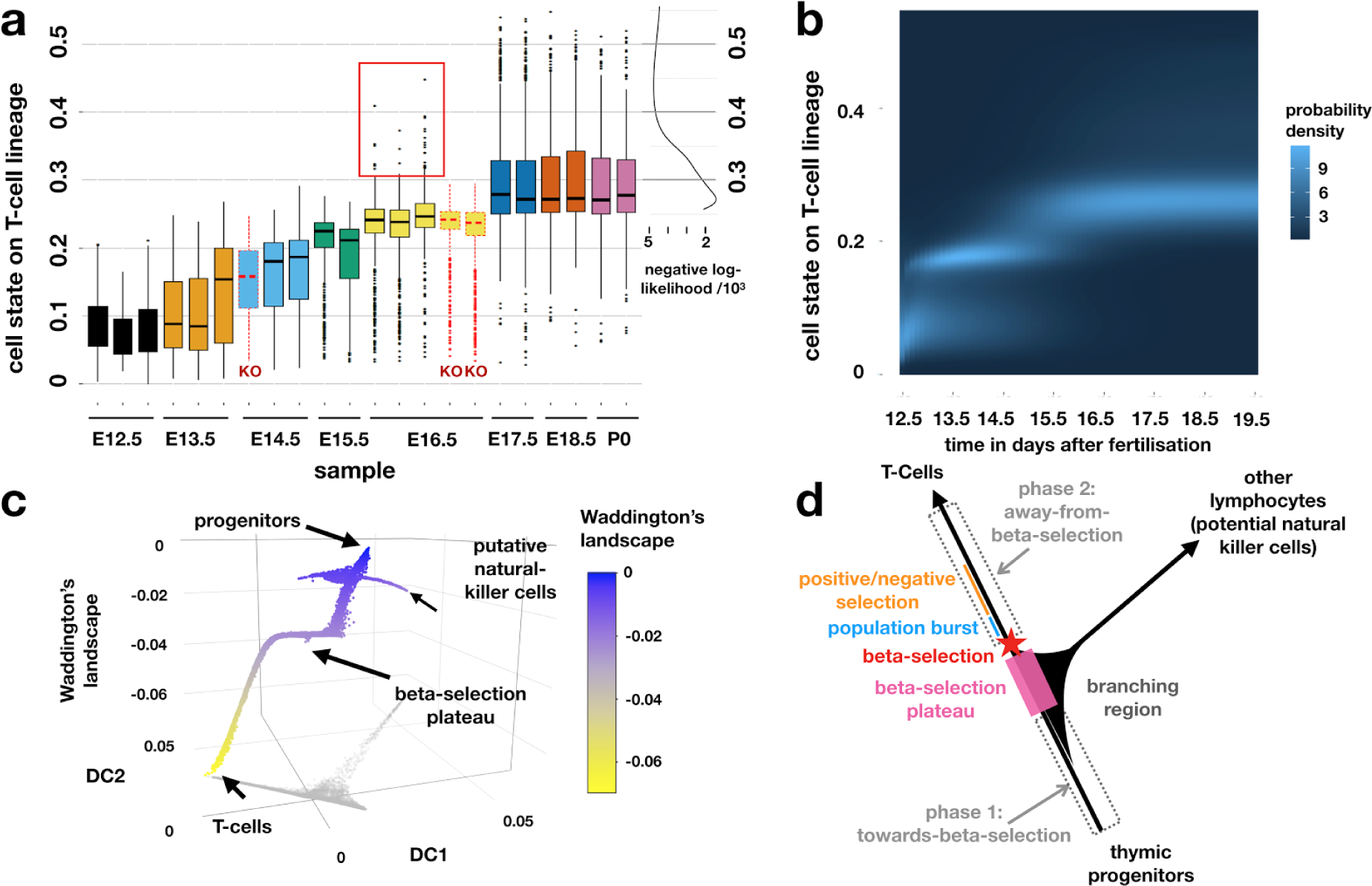
Pseudodynamics annotates a diffusion map of T-cell development with developmental potential, developmental phases and the position of a developmental checkpoint. (**a**) Boxplots of population density in cell state on T-cell lineage by sample with likelihood profile of proposed beta-selection point as function of cell state. The red box denotes high cell state outliers at E16.5 only observed in wild-type. Colour: time point, solid outline: Rag2-KO mice, dotted outline: wild-type mice. (**b**) Heatmap of simulated density on T-cell branch as function of time and cell state. (**c**) Approximation of Waddington’s landscape of the proposed model of thymic T-cell development. (**d**) Schematic model for thymic T-cell development based on pseudodynamics fits.

## ACKNOWLEDGEMENTS

We would like to express our gratitude towards Ping Xu and Kashfia Neherin for mouse husbandry and tissue preparation.

## CONFLICT OF INTEREST

None declared.

### Data and code availability

The Rag2 knock-out Drop-seq data were uploaded to GEO (*not yet public*). The remaining thymus blood data are avaible at GEO (*not yet public*). The pseudodynamics model code is available through GitHub (https://github.com/theislab/pseudodynamics). The visualisation interface for the T-cell data set is avaible at (*not yet public*).

### Supplementary Files

Supplementary Methods Supplementary Figures

### Online Methods Pseudodynamics model

See also (Supp. methods section 2). Pseudodynamics provides a dynamic (time-resolved) description of a population as a density over a cell state coordinate. Development can therefore be modelled as the change in this density over time. In pseudodynamics, the change of this density over time is modelled by a reaction-diffusion-advection model for a non-normalized density across a 1D transcriptomic progression (“cell state”) coordinate (eq. 1). The partial differential equation has a diffusion term with a diffusion parameter field *D*(*s*), dependent on cell state *s*, a drift (or advection) term with a drift parameter *v*(*s*) and a reaction term (the population growth term with a growth parameter *g*(*s*)). All parameters are functions of the cell state *s* and are parameterised as natural cubic splines. The population is the density *u*(*s*, *t*) which gives the relative population size at progression point *s* and time *t*:

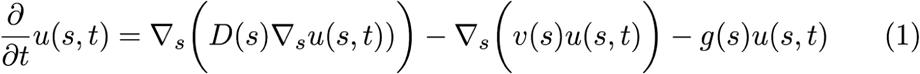

The boundary conditions for this linear system are:

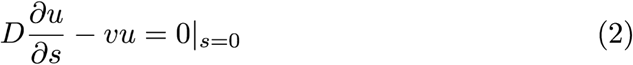

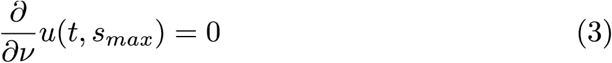

Accordingly, one can formulate a pseudodynamics model for a branching process as system of two coupled partial differential equations. The first partial differential equation describes the evolution of the population along the main trajectory starting at the progenitor state (eq. 4) and the second equation describes the evolution along the side branch starting at the branching region (eq. 5) (Supp. methods Fig. 3). Both equations are coupled at the branching region in which cells can switch between main and side branch.

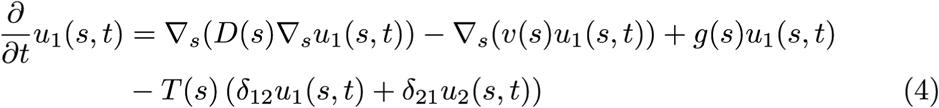

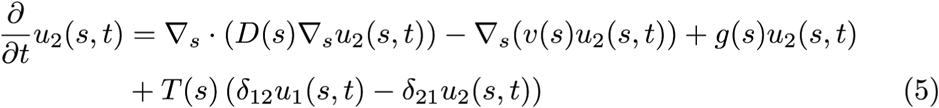

in which the function *T* (*s*) defines a branching region in state space. *T* (*s*) is one inside the branching region and zero outside of it. The branching region is an interval in cell state space with deterministic switching rate δ_*ij*_ from branch *i* to branch *j*. The boundary conditions for this branching model are:

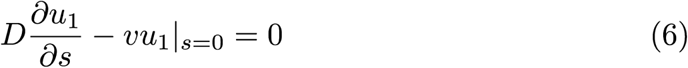

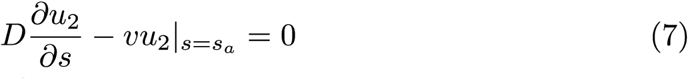

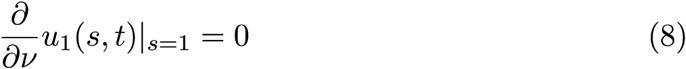

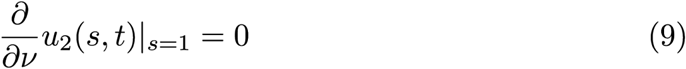

i.e. no-flux conditions for diffusion-advection processes on the left-hand-side and no-flux conditions for diffusion equation on the right-hand boundary. As the advection is artificially decreased to zero at the right boundary, over all no cell mass is lost.

The population size N is a function of time and is defined as the integral of the non-normalized density with respect to cell state:

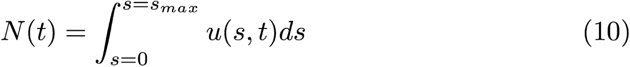

This integral is initialised as the observed population size at the first time point.

### Regularized likelihood and parameter estimation

See also (Supp. methods section 3). The parameters of the pseudodynamics model are estimated from a given dataset using a likelihood *L* (eq. 11) that accounts for the normalized density of the population across the cell state coordinate and the population size. The input data consist of sets of empirical cumulative densities of cell state observation *ecdf*(*s_t,b_*) (pseudotime coordinate) per time point *t* and branch *b*, a set of observed population sizes *N* with corresponding time points and of the standard error in the population size observations per time point 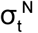:

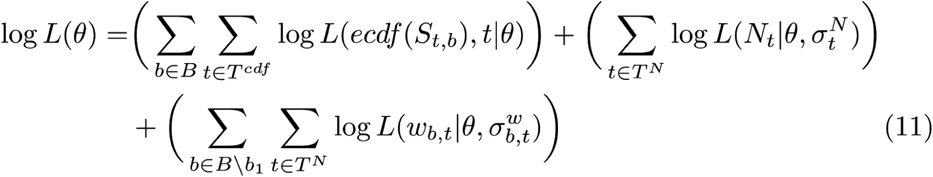

where *B* is the set of branches modelled, *w_b,t_* is the fraction of cells observed in branch *b* at time *t* and 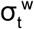 its standard deviation across replicates, *T^cdf^* is the set of time points at which the population density was observed, *s_t,b_* is the vector of cell state observations at time point *t* in branch *b*, *T^N^* is the set of time points at which the population size was observed, *N_t_* is the observed population size at time point *t* and θ is the set of parameters of the pseudodynamics model. Note that the likelihood term on the fraction of cells per branch is not evaluated on the first branch as the proportions across all branches is normalized. The regularized likelihood function *J*, which is maximized for the parameter estimation, is the sum of the negative log-likelihood and smoothness regularisation terms on the parameter splines:

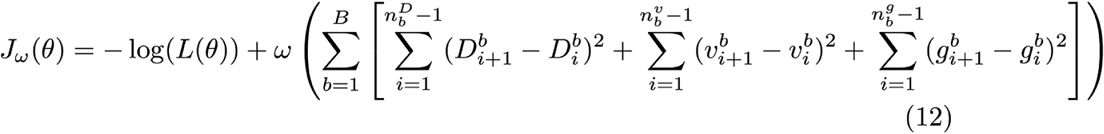

In which ω is the weight of the regularisation terms and 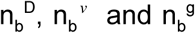 are the number of nodes of the natural cubic splines used to model the parameters *D*, *v* and *g* on each branch *b* out of the *B* branches. The regularization parameter ω was chosen via cross-validation (Supp. methods sec. 4.4.1, Supp. Fig. 13).

The likelihood of the normalized observed population densities in cell state space given a set of parameters is evaluated based on the area between the empirical cumulative density functions (ECDF) of the observed data and simulated cumulative density function. The simulated cumulative density function on branch *b* is:

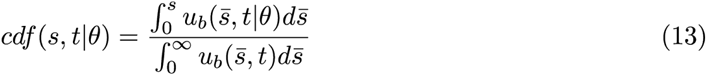

where *u_b_* is the simulated density on branch *b*. We assumed that area between the curves (*A*) is normally distributed with standard deviation σ*^A^*(*t*) and mean σ*^A^*(*t*) estimated based per time point *t* on the area between the curves of the ECDF of each experimental replicates to the ECDF of the union of all cells in all branches of a given time point *S* _t_, yielding the likelihood function

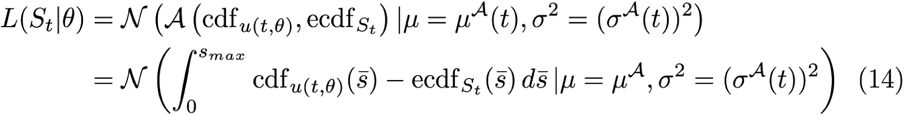

We assumed that the error of the observation of the population size is normally distributed. We estimated the standard deviation of the measurement noise per time point as the standard error of the population size observation at that time point 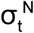. Accordingly, the likelihood for the population size observations is a normal distribution:

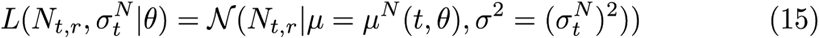

which has the square of the standard error of the population size observations as variance and which has the integral of the simulated density over cell state as a mean parameter:

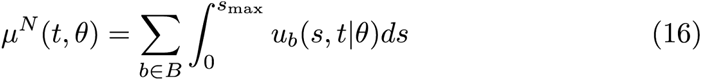

### Implementation of the parameter estimation

See also (Supp. methods section 4). Calibrating the parameters of the pseudodynamics models is non trivial, because for each likelihood evaluation we have to simulate a PDE model. The numerical implementation of the forward simulation of the pseudodynamics model was based on the method of lines. The model was discretized in space using finite volumes. For the solution of the resulting system of ordinary differential equations we employed the Sundials CVODE suite^20^ and AMICI^21^ (https://github.com/ICB-DCM/AMICI/) as a Matlab interface. For the optimization we used a multi-start approach (with gradient information) that is implemented in the Matlab toolbox PESTO^22^ (https://github.com/ICB-DCM/PESTO). We supplied the optimizer with the analytical gradient as this increases efficiency in comparison to numerically computed gradients^23^. Uncertainty analysis and computation of confidence intervals was performed using profile likelihoods (also implemented in PESTO). To determine the regularization parameter, we performed leave-one-out cross validation successively leaving out the data corresponding to a time point and estimating the parameters for the reduced data set. For these parameters we could evaluate the likelihood on the whole data set and compare prediction accuracy.

### Estimation of the cell state coordinate of beta-selection with pseudodynamics

See also (Supp. methods section 5). To compute the likelihood profile of the point of beta-selection across the cell state coordinate s, we trained the pseudodynamics model on the wild type data subset of the combined wild type and knock-out samples diffusion pseudotime model. We adjusted this model for a developmental arrest at a proposed cell state s’. We then estimated the point of beta-selection as a transcriptomic coordinate as the maximum likelihood estimator s* of s’ on the mutant data (eq. 16). Specifically, we set drift parameter at the cell state coordinates beyond the proposed point of arrest to zero (eq. 18,19) and the growth parameter to −3 a lower bound of estimated birth-death parameters. We computed the likelihood profile of s’ between the smallest cell state grid point not observed at the initial time point and the highest cell state observed on the T-cell lineage. We used a least squares objective as a likelihood function to evaluate the fit (eq. 20).

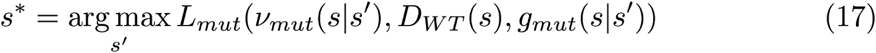

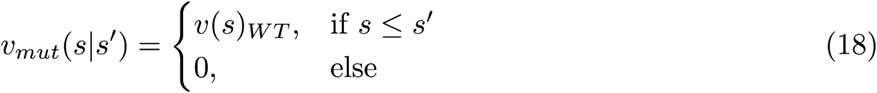

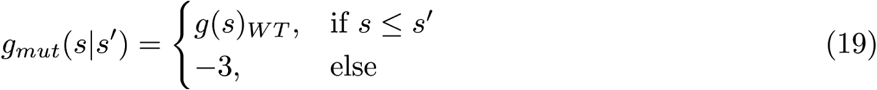

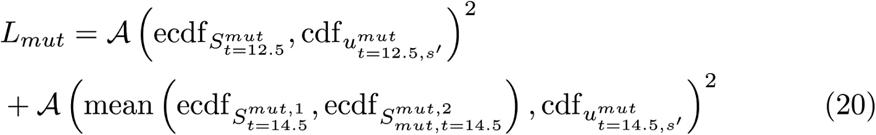

Note that the likelihood profile is a function of the cell state derived from diffusion pseudotime on the union of all wild-type and mutant cells. This diffusion pseudotime coordinate (*s_WT+KO_*) is different from the diffusion pseudotime computed only on the set of wild-type cells (*s_WT_*). We mapped the cell state *s_WT+KO_* back to *s_WT_* to interpret the beta-selection point in the context of the T-cell maturation description established based on the wild-type data (Fig. 2). We note that *s_WT+KO_* is a monotonously increasing function of *s_WT_*. Accordingly, we performed the mapping with a smooth function class (degree 5 natural cubic splines) (Supp. Fig. 17b).

### Computation of Waddington’s landscape

Waddington’s landscape *W* is a developmental potential function of the cell state *s*. We assumed that the gradient of Waddington’s landscape with respect to cell state can be approximated by the drift parameter estimate of the pseudodynamics model (eq. 20). Accordingly, one can approximate Waddington’s landscape as the integral of the negative drift parameter trajectory with respect to cell state (eq. 21s).

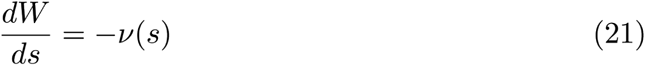

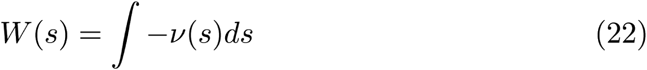

We approximated the integral (eq. 22) with Euler’s method by setting W(0) = 0 and by using the negative drift parameter fit to do step-wise finite difference approximation of W in s (eq. 22), where Δs is the grid spacing of the drift parameter fit in cell state. We initialised W on the second branch with the computed value of W on the first branch at the first cell state of the branching region.

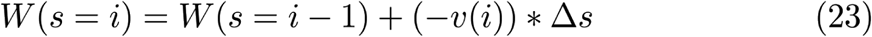

We note that this approach to approximate Waddington’s landscape only yields approximations along the observed developmental trajectories.

### Generation of Drop-seq dataset of T cell development

Detailed description of isolation of thymus resident cells and generation of Drop-seq datasets will be described elsewhere (Kernfeld et al, manuscript in preparation), and raw data files will be available through GEO. Briefly, C57BL6/J and Rag1−/− mice were obtained from The Jackson Laboratory, and thymus tissue isolated from timed pregnant mice. Life cells were enriched using FACS and immediately processed for Drop-seq analysis. Drop-seq was performed following the online protocol provided from the McCarroll lab at Harvard Medical School (Drop-seq Laboratory Protocol version 3.1; http://mccarrolllab.com/dropseq/). Refer to Supp. Fig. 16 for global sample characteristics.

### Drop-seq data processing and analysis

Drop-seq libraries were sequenced at paired-end (20-50) on a Nextseq500. The STAR aligner (v2.4.2), Picard tools (v1.96) and the Drop-seq tools (v1.0) were used to convert raw FASTQ files into digital gene expression matrices. Below, we list the commands and parameters that were used.

| Purpose | Software Package | Command name | Free Parameters |
| --- | --- | --- | --- |
| Convert to SAM | Picard tools | FastqToSam.jar | -- |
| Tag cell barcode | DropSeq tools | TagBamWithReadSequenceExtended | BASE_QUALITY=10, BARCODED_READ=1, NUM_BASES_BELOW_QUALITY=1, DISCARD_READ=False |
| Tag molecular barcode | DropSeq tools | TagBamWithReadSequenceExtended | BASE_QUALITY=10, BARCODED_READ=1, NUM_BASES_BELOW_QUALITY=1, DISCARD_READ=True |
| Remove low-quality barcodes | DropSeq tools | FilterBAM | -- |
| Remove polyA tail | DropSeq tools | PolyATrimmer | MISMATCHES=0, NUM_BASES=6 |
| Convert to FASTQ | Picard tools | SamToFastq.jar | -- |
| Align reads to exome | STAR | -- | -- |
| Sort BAM to speed merging | Picard tools | SortSam.jar | -- |
| Merge with tagging and alignment info | Picard tools | MergeBamAlignment.jar | INCLUDE_SECONDARY_ALIGNMENTS=False |
| Label read with exon | DropSeq tools | TagReadWithGeneExon | -- |
| Screen for bead errors | DropSeq tools | DetectBeadSynthesisErrors | NUM_BARCODES=2000 |
| Form DGE matrix | DropSeq tools | DigitalExpression | MIN_NUM_GENES_PER_CELL=1000 |

We rescaled raw molecule counts of each cell to sum to 10,000, and we transformed the resulting values via X→log2(1+X). The number 10,000 was chosen by rounding the median unique molecular identifier (UMI) count up to the nearest power of 10.

T-cell receptor alignment was improved by augmenting the reference genome. The augmented reference contained an artificial TCR contig in which known constant, joining, and variable regions of the TCR were concatenated. TCR regions were extracted from TRACER annotation files. The boundaries were annotated as splice junctions, allowing the extensive spliced alignment capabilities of STAR to position reads despite TCR rearrangement. Reads aligning to the TCR contig were subsetted, and transcript quantification was performed as above. We made two alterations: in place of MIN_NUM_GENES_PER_CELL=1000, we used cell barcodes established using the conventional alignment pipeline, and we specified READ_MQ=1. TCR realignment was performed after the initial analysis, and it did not affect the set of cells classified as blood.

### *In silico* isolation of wild-type blood cells E12.5-E16.5

Quality control and blood isolation was conducted using the R language (https://www.R-project.org/) and the package ‘Seurat’^24^ (http://www.satijalab.org/seurat). The main goals for quality control were to verify exclusion of female embryos; to exclude empty droplets; and to deplete cell doublets. Only male embryos were analyzed to avoid biological confounding by sex. To remove empty droplets, we excluded any cell expressing less than 1000 genes. We also excluded any gene expressed in less than 10 cells. Doublet depletion was carried out, followed by isolation of the blood. Both steps used unsupervised machine learning.

For doublet depletion and blood isolation, we used two pipelines that differ only in their final steps. Each began by compensating for variation due to the cell cycle. For each of five cell cycle phases (IG1.S, S, G2.M, M, M.G1), scores were computed by averaging expression within each cell over a set of genes found in the second workbook of table S2 from a reference^25^. Seurat’s RegressOut function was used to replace expression levels with standardized residuals from linear regressions (one per gene). In each regression, observations are cells; the response variable is log-normalized expression; and the covariates are the five cell cycle scores. After cell-cycle correction, we enriched for informative genes by applying Seurat’s MeanVarPlot function (with x.low.cutoff = 0.1 and y.cutoff = 0.5). Principal components analysis (PCA) was run on the selected genes using as features the normalized residuals from RegressOut.

The first difference in the two pipelines occurs after PCA. For doublet removal, the top 20 principal components (PCs) were used as input to Barnes-Hut t-Stochastic Neighbor Embedding (tSNE). DBSCAN was used to isolate and remove outlying cells and clusters showing markers from multiple cell types. In DBSCAN, the tSNE embedding was used as input, and the parameters were 1.1 (neighborhood size) and 5 (minPts). In total, 52 putative doublets and 80 outlying cells were excluded from downstream analyses.

For isolation of the blood, the entire process up to PCA was repeated after doublet depletion. Clustering was then carried out using Seurat’s FindClusters function, which applies a variant of the Louvain algorithm^26^ to a shared-nearest-neighbor graph constructed in the principal subspace (20 PCs, resolution 0.5). Results were visualized as before via tSNE. Six contiguous clusters were manually labeled as blood based on expression of known markers. Different parameter choices for variable gene selection and for the number of principal components were explored and results remained qualitatively consistent.

### *In silico* isolation of E17.5-P0 wild-type blood cells and E14.5 *Rag2*−/− blood cells

The blood from these later time points was aligned following the same procedure. Data were filtered for at least 1000 genes per cell, but the requirement was loosened to at least 3 cells per gene due to the smaller total number of cells. No doublet removal was attempted. Blood isolation was performed via the pipeline described above using x.low.cutoff = 0.1 and y.cutoff = 1.2 for gene selection, 25 principal components, and the Louvain algorithm with resolution 0.5. Two clusters lacked *Ptprc* and expressed thymic stromal markers, and these were manually removed. Different parameter choices were explored for variable gene selection and for the number of principal components; relabeled results remained relatively robust. Wildtype cells were processed together and *Rag2-/-* cells were processed separately.

### *In silico* isolation of E16.5 *Rag1*−/− blood cells

For whole-thymus samples from *Rag1* mutant embryos, the same alignment, quantification, and quality control steps were performed (>1000 genes per cell, >3 cells per gene). Cells were classified by k-nearest-neighbors (k=25) after projection into a 20-dimensional principal subspace, with both PCA and classifier trained on the E12.5-E16.5 wt data. Cells classified into any of the six blood clusters were retained for analysis.

### Preparation of pseudodynamics input from blood cell transcriptomes

(See also Supp. methods sec. 6.2.5). We fit diffusion pseudotime with one branching point to the union of all lymphocytes from all samples with scanpy^27^ (k=100, knn=False) (diffusion map A). We classified the resulting four groups of cells based on markers genes as progenitors/intermediate cells (Flt3), T-cells (Cd8b1, Cd4), putative natural killer cells (Klrk1, Ifng) and putative dendritic cells (H2-Aa, H2-Ab1, H2-Eb1) (Supp. Fig. 2). We discarded the putative dendritic cell group to obtain the data set that contains a single branching between the αβ-T-cell lineage and a lineage of other lymphocytes (putative natural killer cell lineage) and fit a new pseudotemporal ordering on this data set with scanpy^27^ (k=100, knn=False) (diffusion map B). We defined the allocations of cells to branches and branching region in diffusion map B based on pseudotime coordinates and diffusion component 1 and 2 coordinates (Supp. methods). We repeated the workflow from diffusion map A to diffusion map B separately for the wild-type only and the wild-type with knock-out samples data sets. We discarded the dendritic cell group and the natural killer cell group from diffusion map A to obtain the data set that contains no branching and only the T-cell lineage (used for monocle2-based cell state coordinates^5^).

